# Modulation of sensory input to the spinal cord by peripheral afferent fibres. Searching for relays

**DOI:** 10.1101/2024.12.04.626763

**Authors:** Urszula Sławińska, Ingela Hammar, Elzbieta Jankowska

**Affiliations:** Department of Neuroscience and Physiology, Sahlgrenska Academy, University of Gothenburg, Sweden; Nencki Institute of Experimental Biology PAS, Poland

**Keywords:** Myelinated nerve fibres, excitability, astrocytes, GABA, rat

## Abstract

A long-lasting GABA-dependent increase in the excitability of afferent fibres, and thus modulation of the sensory input to the spinal cord, may be evoked by epidural polarization. However, the direct effects of fibre polarization are short-lasting and the sustained increase in their excitability appears to be secondary to the release of GABA from nearby astrocytes. We have now investigated whether the modulation of spinal sensory input by stimulation of a peripheral nerve, previously attributed to synaptically evoked intraspinal field potentials, is evoked in a similar way. However, as neither its dependence on GABA nor its relays have been investigated, we addressed the question of whether the increase in the excitability of epidurally stimulated afferent fibres following a peripheral nerve stimulation does or does not depend on GABA and whether it might be mediated by astrocytes. The effects of conditioning stimulation of the tibial nerve were evaluated from changes in the excitability of both group I and group II muscle afferents, estimated from action potentials recorded in peripheral nerves and in field potentials recorded in the dorsal horn respectively in acute experiments on deeply anaesthetized rats. The excitability of the afferents was increased by stimulation of group II and/or cutaneous but not group I muscle afferents. The effects were significantly weakened by blocking GABA channels by gabazine, and by astrocyte toxin L-alpha-aminoadipic acid (L-AAA), indicating that the excitability of both group I and group II afferent fibres may be modulated by GABAergic astrocytes, the new role played by astrocytes.

## INTRODUCTION

Input to the spinal cord may be modulated at several sites, between the origin of the sensory information in peripheral receptors to the synaptic contacts between afferent fibres and their spinal target cells. We have recently focused on one of these sites, where afferent fibres branch profusely at the border between the dorsal column and the dorsal horn and where the polarization of the dorsal columns evokes a long-lasting increase in the excitability of these fibres (Jankowska *et al*., 2017; Li *et al*., 2020; Jankowska *et al*., 2022; Hammar & Jankowska, 2024). The polarization-evoked sustained increase in the excitability of afferent fibres within their branching region was found to depend on GABA (Li *et al*., 2020), most likely acting on extrasynaptic GABA receptors distributed in close vicinity to Na^+^ channels (Lucas-Osma *et al*., 2018; Hari *et al*., 2022).

GABAergic interneurons were originally considered as a source of GABA increasing the excitability of afferent fibres within their proximal branching region (Hari *et al*., 2022). However, recent observations indicate that the increase in the excitability evoked by epidural polarization greatly depends on GABAergic glial cells, in particular astrocytes, because it is prevented by astrocyte toxin L-alpha-aminoadipic acid (L-AAA)(Hammar & Jankowska, 2024). Astrocytes have already been shown to increase fibre excitability in the visual cortex by affecting Nav1.6 ion channels (Ryczko *et al*., 2021) and in the trigeminal nucleus by lowering [Ca2^+^] “at focal points along the axons” (Gaudel *et al.,* 2024). Astrocytes may thus be involved in modulating not only the synaptic transmission between neurons (for review see e.g. Fellin, 2009; Allen, 2014; Hanani & Verkhratsky, 2021) but also the input to the neurons at a stage before it reaches them.

Previous observations have demonstrated how some GABA-mediated modulatory actions of astrocytes may be evoked. In particular, the study of the Christensen group (Christensen *et al*., 2018) revealed not only that 29% of dorsal horn astrocytes in the turtle are GABAergic but also that they may release GABA. Hence, they may evoke a long-lasting increase in the concentration of GABA in the area of afferent branching in the dorsal columns.

The main aim of the present study has been to find out whether modulation of spinal afferent input by stimulation of peripheral afferent fibres (Bączyk & Jankowska, 2018) is GABA-evoked and, if so, if it may be mediated by astrocytes in a way similar to that following epidural polarization. Bączyk & Jankowska (2018) demonstrated that stimulation of a mixed hindlimb nerve is followed by an increase in the excitability of epidurally stimulated fibres. This was considered as an effect of extracellular field potentials evoked in the dorsal horn, replicating the effects of field potentials evoked by epidural polarization. However, the modulation of the excitability of afferent fibres evoked in this way might likewise be mediated by astrocytes releasing GABA. If so, the facilitatory effects of the conditioning stimulation of a peripheral nerve would be weakened or prevented by astrocyte toxin L-AAA and the GABA channels blocker gabazine.

Provided that astrocytes modulate the excitability of afferent fibres, the second aim of the study was to estimate how specific their actions are. In particular, whether they contribute to the depolarization of not only proximal parts of the afferents but also enhance or counteract the presynaptic inhibition and/or PAD (primary afferent depolarization) evoked via axo-axonic contacts of GABAergic interneurons within their distal compartments. Proximally and distally distributed changes in the excitability of afferent fibres were considered to be mediated by subpopulations of GABAergic interneurons at different locations (Hari *et al*., 2022; Lin *et al*., 2023; Metz *et al*., 2023) (see Figs. 3 and 7 in Hari *et al*., 2022) with only their ventral subpopulation mediating the specific patterns of presynaptic inhibition and PAD. However, proximally and distally evoked depolarization of afferent fibres might also be mediated by subpopulations of astrocytes or by astrocytes and GABAergic interneurons respectively. We, therefore, evaluated effects of L-AAA on changes attributable to presynaptic inhibition evoked by stimulation of peripheral afferents. This was done by comparing changes in monosynaptic components of extracellular field potentials recorded in the dorsal horn by stimulation of a peripheral nerve and by epidural stimuli. Similar depression of both was expected to be compatible with their common mechanism while differences would indicate distinct actions of GABAergic astrocytes and interneurons.

## METHODS

All experiments were approved by the Regional Ethics Committee for Animal Research (Göteborgs Djurförsöksetiska Nämnd, permit no 5.8.18-16183/2019) and followed EU and NIH guidelines for animal care. The animals (19 female adult Sprague Dawley rats, 230-350 g, Janvier Labs, France) were housed under veterinary supervision at the Laboratory of Experimental Biomedicine at Sahlgrenska Academy with food and water ad libitum. Measures were taken to minimize animal discomfort by preceding the intraperitoneally applied anaesthetics with a short period of isofluorane inhalation anaesthesia when the animal was unrestrained in a box. The number of animals was minimized by using protocols that allowed more than one question to be addressed in each experiment. The experimental procedures were as described by Hammar & Jankowska (2024).

### Design of the experiments

The main elements of the experimental design are illustrated in Figure 1a. The circle at the border between the dorsal column and the dorsal horn indicates the site where the excitability of afferent fibres providing input to the spinal cord was investigated, using effects of epidural stimulation of these fibres as a measure. Their excitability was estimated from the size of the compound action potentials recorded from the common peroneal (Per) nerve (referred to as “nerve volley”) and the size of extracellular field potentials evoked in the dorsal horn (referred to as “field potential”), as indicated in Figure 1b. The epidural stimulation was applied within the segment where the largest afferent volleys and cord dorsum potentials were evoked from the peroneus nerve and where it is most potently modulated by epidural polarization and astrocytes (see Hammar & Jankowska, 2024). The dorsal horn field potentials were recorded at a location where the largest field potentials were evoked by group II but not group I Per muscle afferents.

**FIGURE 1.**
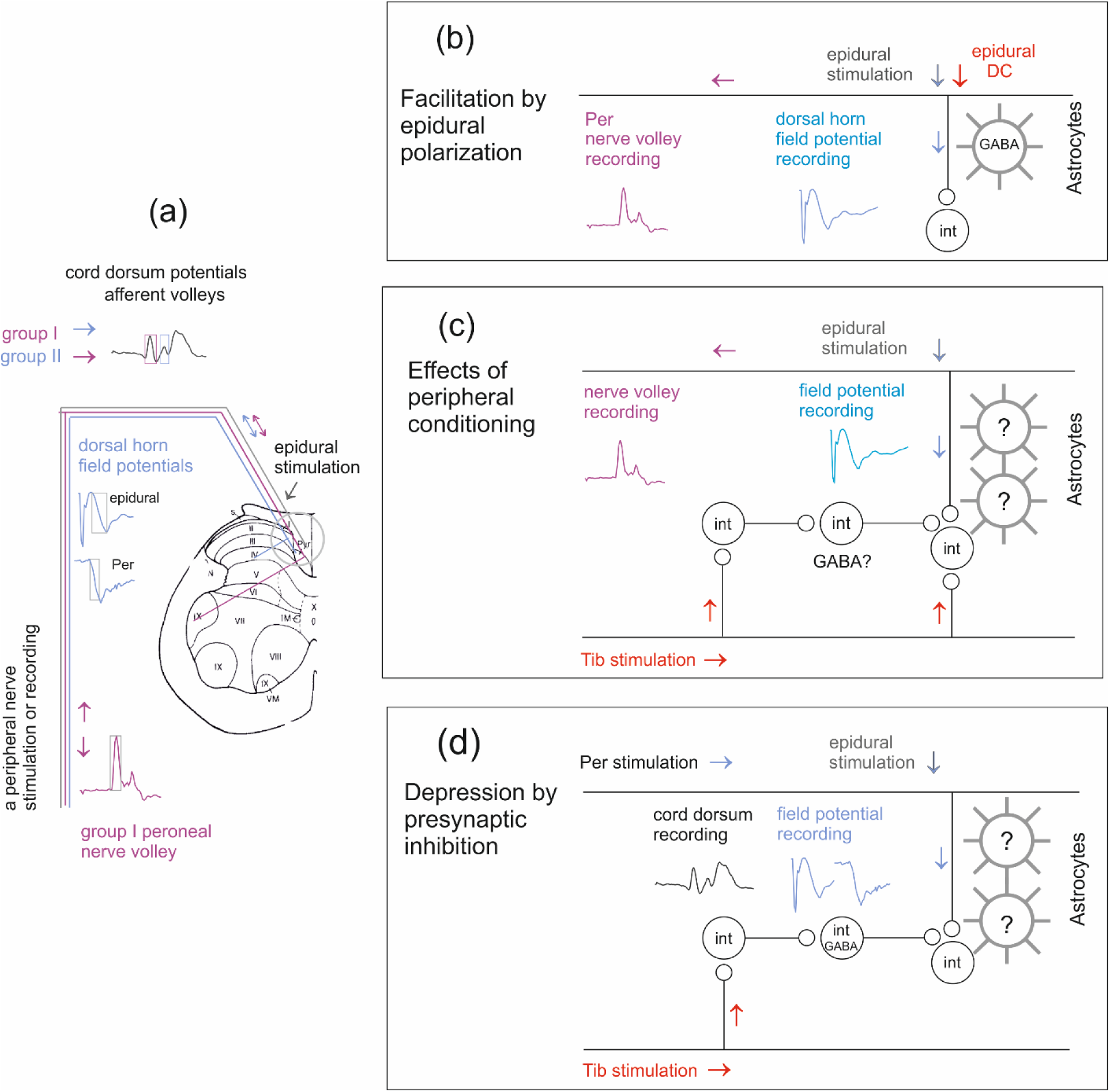
Diagrams of experimental paradigms. (a) The preparation used in all the experimental series in which afferent fibres traversing the dorsal columns at the L3-L4 lumbar level were stimulated epidurally. The fibres included group I (purple), group II (blue) muscle afferents and skin afferents (grey) but activation of group I and group II muscle afferents was differentiated based on their specific responses: the shortest latency compound action potentials attributable to group I afferents in a peripheral nerve (Per) and the earliest components of field potentials reflecting monosynaptic excitation of neurons within the explored region of the dorsal horn by group II but not group I afferent fibres. The sample records illustrate the responses used for analysis. The circle indicates the region within which the changes in the excitability were expected to occur. (b) and (c) Experimental paradigms for testing changes in the excitability of epidurally stimulated afferent fibres evoked by epidural polarization reported previously (Hammar & Jankowska, 2024) and by conditioning stimulation of the tibial nerve. Blue and red arrows indicate the direction of action potentials induced by the test and conditioning stimuli. The time windows within which the responses evoked by these stimuli were compared are indicated by grey rectangles on the sample records in (a) and Figure 2. (d) Experimental paradigm for testing effects of the same conditioning stimuli on potentially occurring presynaptic inhibition of transmission from epidurally stimulated fibres. Astrocytes postulated to mediate changes in the excitability of the analysed fibres within their branching regions are indicated to the right. Relay neurons are indicated as circles without specifying the types of synaptic contacts that they form. For further explanations see the text.

Modulation of the excitability of epidurally stimulated fibres was evoked by conditioning stimulation of the tibial (Tib) nerve, containing muscle and skin afferent fibres. The nerve was routinely stimulated at an intensity near maximal for group II muscle spindle afferents (four to five times the threshold) but the comparison of effects of stimuli was done using different intensities.

Only the earliest components of the nerve volleys and field potentials, within time windows of 0.4-0.9 ms and thus compatible with directly evoked potentials were taken into account. Their increase indicated an increase in the number of afferent fibres excited by epidural stimulation following conditioning stimulation, which in turn indicated an increase in the excitability of these fibres. The effects of the test and conditioning stimuli were compared before and after local administration of the astrocyte toxin L-AAA and the GABAA channel blocker gabazine. L-AAA and gabazine were administered by diffusion from a micropipette in contact with the surface of the dorsal column within a 1 mm distance from the tip of the electrode used for epidural stimulation and compared to the effects of epidural polarization, as indicated in Figure 1c.

The degree of presynaptic inhibition was estimated from the decrease in the dorsal horn field potentials evoked from Per that followed the conditioning stimulation of Tib as indicated in Figure 1d, at conditioning-testing intervals of 19-22 ms.

### Preparation

Anaesthesia was induced with isoflurane (Baxter Medical AB, Kista, Sweden; 4% in air) followed by intraperitoneal administration of pentobarbital sodium (Apoteksbolaget, Göteborg, Sweden; 30 mg/kg) together with α-chloralose (Acros Organics, Geel, Belgium, 30 mg/kg). The anaesthesia was supplemented with an intraperitoneal injection of α-chloralose (3 mg/kg, up to 40 mg/kg) at 3-4 hours intervals. When a deep level of anaesthesia had been established, the neuromuscular transmission was blocked during the experiment by gallamine triethiodide (Sigma Aldrich, G8134). Gallamine was injected intravenously (via the tail vein) at an initial dose of 10 mg/kg supplemented with 5 mg/kg when needed, either i.v. or intraperitoneally. Artificial ventilation was applied via a tracheal tube by a respiratory pump (CWE; model SAR-830/P) set to maintain the end-tidal CO2 level at ∼3.4–4.5%. The core body temperature was kept at ∼38°C using servo-controlled heating lamps. To compensate for fluid loss, 10 ml of a glucose bicarbonate buffer (0.84g NaHCO3 and 5g glucose in 100 ml distilled H2O) were injected subcutaneously during the preliminary dissection. Throughout the experiment, the state of the animal was monitored by recording ECG, via needle electrodes inserted subcutaneously over the thorax, the concentration of CO2 in the expired air and the rectal temperature. The experiments were terminated by a lethal injection of anaesthetics resulting in ECG-verified cardiac arrest.

The initial surgery included the dissection of two hindlimb nerves, the mixed peroneal (Per) and tibial (Tib) nerves. These were transected distally, freed up to the mid-hip level and mounted on pairs of silver electrodes in a paraffin oil pool maintained at 32-35°C. The vertebral column was immobilised with clamps, between the Th13 and L1 levels, a laminectomy was performed exposing the L2–L5 spinal segments and the cord covered with paraffin oil maintained at 35-37 °C. The dura remained intact apart from a small opening made for the introduction of the recording glass micropipette and for the drug administration.

### Stimulation and epidural polarisation

Afferent fibres in Per and Tib nerves were stimulated epidurally at the level of the L3-4 segments of the spinal cord, above the Per and Tib motor nuclei where the most extensive branching of group I and II muscle afferent fibres occurs. The stimulation was applied within the 2–3 mm length of the spinal cord over which the largest afferent volleys and cord dorsum potentials were evoked (see Figure 1b in Li *et al*., 2020) from the two nerves at the dorsal root entry zone. The nerves were stimulated bipolarly using constant voltage stimuli at intensities 3–5 times the threshold. Afferents traversing the dorsal columns were stimulated monopolarly by a tungsten needle electrode insulated except for 20–30 μm at the tip (Microneurography active needle, UNA35FNM, FHC, Bowdoin, ME, USA; impedance 70– 400 kΩ) against a subcutaneous abdominal reference electrode. Single 0.2 ms constant current rectangular stimuli were usually delivered at intensities 16-18 μA and were submaximal for both nerve volleys recorded from a peripheral nerve and for field potentials recorded within the dorsal horn, the threshold stimuli for both being similar (10-12 µA). The stimuli were applied epidurally to avoid direct contact between the electrodes and the nervous tissue (see Holsheimer, 2002; Bikson *et al*., 2013; Holsheimer & Buitenweg, 2015; Jackson *et al*., 2017). Epidural polarisation was passed via the same tungsten electrode (see (Bączyk & Jankowska, 2018). It was delivered via a custom-designed polarizer (Magnusson, D, Göteborg University) using 1 μA depolarisation for 1 min.

### Recording

The orthodromic afferent volleys evoked by peripheral nerve stimulation and synaptically recorded field potentials electrotonically spreading to the surface of the spinal cord were recorded close to the dorsal root entry zone by a silver ball electrode touching the dura matter against a reference electrode inserted into a back muscle. Extracellular field potentials evoked by stimulation of either the peripheral nerves or epidurally were recorded with glass micropipettes with a tip of about 2 µm at 0.7 - 0.9 mm depth below the surface of the dorsal column. These were inserted through a small hole in the dura mater about 1 mm rostral to the tip of the tungsten electrode at an angle of 7 deg with the tip directed laterally. Single records, as well as averages of 10 consecutive records, were sampled and stored online (sampling frequency 1 Hz; low passband filter set to 10 or 15 Hz and high-pass band filter at 5 or 3 Hz, resolution of 0.03 ms).

Changes in the size (area) of compound action potentials recorded in peripheral nerves and earliest components of field potentials (within 0.4-0.7 ms time windows) were used as a measure of the number of excited nerve fibres and thus of their excitability. Later components of the field potentials (within 1.5-2.5 ms time windows) were used for the comparison of di- and tri-synaptically evoked components of these potentials.

### Drug application

The drugs were allowed to diffuse from a glass micropipette (tip diameter about 3 µm) in contact with the surface of the dorsal column filled with 1 mM solution of L-alpha-aminoadipic acid (L-AAA), A7275 Sigma, or 100 mM solution of gabazine, SR-95531 Merck.. The diffusion of L-AAA and gabazine started 15-30 min and 10-15 min prior to the beginning of the recording respectively and continued during the whole period of recording (for 30-90 min). The effects of L-AAA and gabazine were estimated only in preparations in which they modified the size of the control nerve volleys or field potentials, indicating that the experimental conditions were adequate.

### Analysis

The areas of nerve volleys were estimated using a custom-designed analysis programme (designed by E. Eide, Göteborg University, see Jankowska *et al*., 1997). They were expressed in arbitrary units in % of control records. The areas of the earliest components of the nerve volleys and field potential evoked by epidural stimulation were measured within a time window of 0.4-0.7 ms from their onset (see boxed areas in sample records in Figure 1a). The time windows used for measurements of field potentials evoked by stimulation of peripheral nerves were somewhat longer (0.5-0.9 ms) taking into account less synchronous actions of potentials reaching their spinal targets because of the longer conductance distance. The epidurally evoked nerve volleys were evoked at latencies corresponding to the latencies of afferent volleys in group I muscle afferents indicating that they were evoked in nerve fibres with the same conduction velocity, i.e. likewise group I afferents.

Student’s *T*-test was used to evaluate differences between means (± SD) of normally distributed experimental data (Shapiro-Wilk test or D’Agostini and Pearson test for the selected pre- and post-conditioning, pre- and post-L-AAA, and pre- and post-gabazine data. In the case of lack of data normal distribution, the nonparametric Kruskal-Wallis test with Dunn’s post hoc method for multiple comparisons was used (∗ *p* < 0.05; ∗∗ *p* < 0.01; ∗∗∗ *p* < 0.001; ∗∗∗∗ *p* < 0.0001). The statistical analysis was performed using GraphPad Prism 10.0.3.

## RESULTS

### Comparing changes in the excitability of group I and II muscle afferent fibres at their entry to the spinal cord following conditioning stimulation of peripheral afferents

The excitability of epidurally stimulated *group I muscle afferent fibres* was estimated from the size of the earliest components of the compound action potentials recorded from Per, as indicated in Figure 1a. They fulfilled the requirements of action potentials evoked in group I afferents because they were evoked at the same latency as the latency of anterogradely conducted afferent volleys recorded at the level of the dorsal root entry zone following stimulation of the lowest threshold muscle afferents (with examples in Figure 3), indicating the same conduction time as between the peripheral nerve and the spinal cord. The size of these nerve volleys was used as a measure of the number of the excited group I afferent fibres, with an increase following stimuli of the same intensity taken to indicate increased excitability of the epidurally stimulated fibres.

The excitability of *group II muscle afferents* was estimated from the size of extracellular field potentials evoked in the dorsal horn using a time window for their monosynaptic components as indicated in Figure 1a. The field potentials from Per were recorded at a location (depth 0.7-0.9 mm from the surface of the dorsal column) where field potentials from group I afferents were absent and where both extracellular field potentials and neuronal activity were evoked at stimulus intensities exceeding those near-maximal for group I afferents and followed the group II components of afferent volleys with 0.3-0.5 ms delays, as illustrated in the rightmost expanded records in Figure 2. These field potentials were therefore attributed to group II muscle afferents. Maximal field potentials were usually evoked at 4 or 5T stimulus intensities, although larger field potentials were sometimes evoked by stronger (up to 10T) stimuli. Whether this depended on differences in the proportions of skin afferents or topographic factors (more medially, laterally or rostrocaudally located recording sites) has not been determined. Neither was the origin of small field potentials preceding those attributed to group II afferents defined but they might have been evoked within a neighbouring region by group I afferents or skin afferents. The records of field potentials were supplemented by records from cord dorsum potentials (lower records in Figure 2) reflecting the intraspinal field potentials attributable to group II afferents.

**FIGURE 2.**
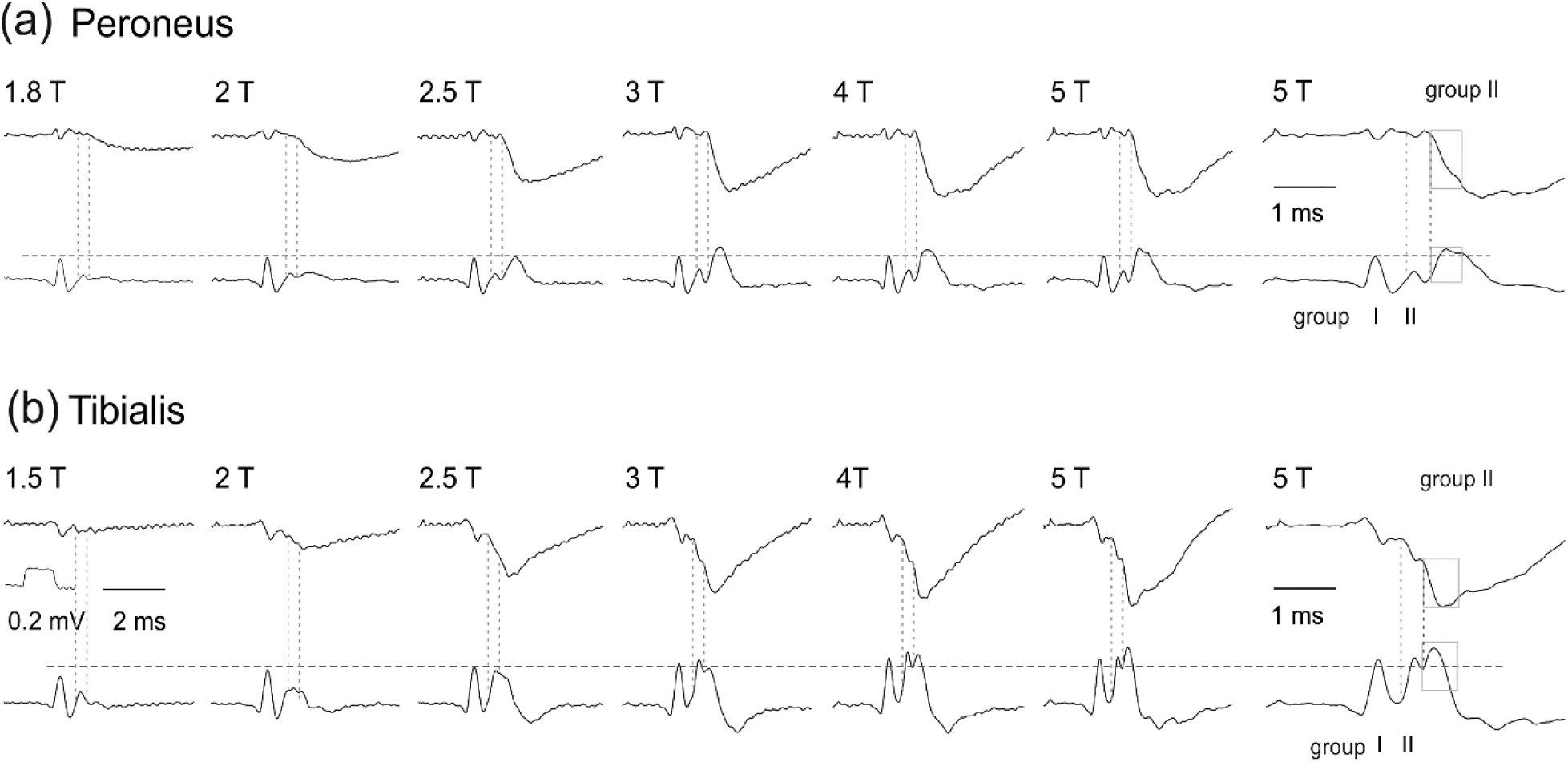
Origin of field potentials within the explored region of the dorsal horn. Upper records, extracellular dorsal horn field potentials evoked by graded stimulation of (a) Per and (b)Tib nerves, recorded at a depth of 0.85 mm. Lower traces, afferent volleys and cord dorsum potentials evoked simultaneously with the field potentials, recorded from the surface of the spinal cord at the dorsal root entry zone. The vertical dotted lines indicate the estimated onset of afferent volleys evoked by group II muscle afferent fibres and extracellular field potentials evoked by them respectively. Note that the early components of afferent volleys from Per and Tib were near maximal at about 2T and 3T respectively (the level of the horizontal dotted lines and that the field potentials following the second vertical lines grew with stimulus intensities between 2 and 5T in parallel with the second components of the afferent volleys and the cord dorsum potentials. In the rightmost, twice horizontally expanded records, the time windows for measuring the monosynaptically evoked components of the field potentials and cord dorsum potentials are indicated by squares. Note that the first components of field potentials from Tib started too early to be evoked by group II afferents (as they were evoked at the same latency as the group II afferent volleys) and at a threshold too high to allow their attribution to group I afferents. They might thus been evoked by skin afferents. For further indications see text.

Conditioning stimulation of Tib, following the experimental design outlined in Figure 1c, was found to increase the excitability of both group I and group II muscle afferents as it increased the earliest components of nerve volleys following epidural stimulation (Figure 3b) as well as extracellular field potentials evoked in the dorsal horn (Figure 3a). The control records of these potentials (blue or purple) and those following stimulation of Tib (black) are superimposed to ease their comparison. They illustrate the common finding that both were increased at the conditioning stimulation intensities within the range for group II afferents (2-5T). Only a weak increase followed stimuli <2T, i.e. below threshold for Tib group II afferents, indicated by the first dotted vertical lines in Figure 2b. Increasing the intensity of the conditioning stimuli above that maximal for group II muscle afferents (about 4T) had only a minor effect. As the conditioning stimuli at intensities below 2T were not effective, the probability of the contribution of group I muscle afferents was very low. A major contribution of the fastest-conducting skin afferents was also unlikely as a considerable proportion of these afferents would have been excited by 2T stimuli.

The degree of the increase was evaluated when single stimuli rather than trains were used as conditioning stimuli since repetitive stimulation evokes stronger presynaptic inhibition or post-activation depression of synaptically evoked field potentials that might obscure estimates of changes in the excitability of the fibres evoking these potentials. The effects of these stimuli were tested on submaximal responses evoked by epidural stimuli expected to leave a sufficient subthreshold fringe to allow the expression of the effects of conditioning stimuli. As illustrated in Figure 3, the relative differences between the test and conditioned responses were indeed largest for the mid-range intensities of epidural stimulation (d) and mid-sized field potentials evoked by them (e).

Following the maximal conditioning stimulation (4-5T) of Tib, the increase in nerve volleys induced in group I afferent fibres and in the earliest components of the dorsal horn field potentials evoked by group II afferent fibres was found in all 19 experiments. The mean relative increase in nerve volleys and field potentials fell within the same range (155-175% see Table 1) and was not found to be statistically significantly different (ns; *p* > 0.05) whether at 5T or lower intensities of the conditioning stimuli.

**TABLE 1.** Effects of conditioning Tib stimulation under different experimental conditions. *(either the version in colour or the black-white version* Comparison of effects of Tib stimulation on dorsal horn field potentials and nerve volleys recorded from Per, both evoked by epidural stimulation, as well as on presynaptic inhibition of field potentials evoked at the same site by Per stimulation. Standard intensities of the test stimuli were submaximal and of conditioning stimuli near maximal for group II muscle afferents in Tib (5T). The comparison involved the effects of these stimuli under control conditions (column b) and during topical administration of either the astrocyte toxin L-AAA (column c) or gabazine (column d) on the dorsal surface of the spinal cord. The effects of conditioning stimulation of Tib in the presence of L-AAA and gabazine are expressed in % of control responses (columns c and d) and as the difference between them and the effects evoked under control conditions (column e). Column f shows that the differences were all statistically significant except for the effects of Tib conditioning of the presynaptic inhibition following L-AAA. Column g shows that no statistically significant differences were found between the effects of either L-AAA or gabazine on field potentials or nerve volleys (comparisons 1-3 and 2-4). However, both the nerve volleys in group I afferent fibres and presynaptic inhibition of transmission from group II afferent fibres were significantly more affected by gabazine than by L-AAA (comparisons 3-4 and 5-6). Columns h and i show the number of rats in experiments and the number of experimental series in which these data were obtained.

|  | Control | After L-AAA |  | After gabazine |  | difference | P value paired t-test |  | P value unpaired | No of rats | No of series |
| --- | --- | --- | --- | --- | --- | --- | --- | --- | --- | --- | --- |
| a | b | c |  | d |  | e | f |  | g | h | i |
| Facilitation: | 162 ± 49 | 129 ± 40 | 1 |  |  | 31% | 0.006 | 1-2 | 0.8 | 3 | 8 |
| Field potentials (group II) | 164 ± 38 |  |  | 124 ± 27 | 2 | 40% | 0.0004 | 1-3 | 0.61 | 5 | 11 |
| Facilitation: | 175 ± 22 | 136 ± 10 | 3 |  |  | 39% | 0.002 | 3-4 | 0.03 | 4 | 9 |
| Nerve volleys (group I) | 155 ± 49 |  |  | 116 ± 11 | 4 | 39% | 0.04 | 2-4 | 0.38 | 5 | 10 |
| Facilitation: | 80 ± 10 | 81 ± 6 | 5 |  |  | -1% | 0.54 | 5-6 | 0.005 | 4 | 8 |
| Per field potentials (group II) | 85 ± 10 |  |  | 97 ± 8 | 6 | -12% | 0.002 |  |  | 5 | 6 |
|  | Control | After L-AAA |  | After gabazine |  | difference | P value paired t-test |  | P value unpaired | No of rats | No of series |
| a | b | c |  | d |  | e | f |  | g | h | i |
| Facilitation: | 162 ± 49 | 129 ± 40 | 1 |  |  | 31% | 0.006 | 1-2 | 0.8 | 3 | 8 |
| Field potentials (group II) | 164 ± 38 |  |  | 124 ± 27 | 2 | 40% | 0.0004 | 1-3 | 0.61 | 5 | 11 |
| Facilitation: | 175 ± 22 | 136 ± 10 | 3 |  |  | 39% | 0.002 | 3-4 | 0.03 | 4 | 9 |
| Nerve volleys (group I) | 155 ± 49 |  |  | 116 ± 11 | 4 | 39% | 0.04 | 2-4 | 0.38 | 5 | 10 |
| Facilitation: | 80 ± 10 | 81 ± 6 | 5 |  |  | -1% | 0.54 | 5-6 | 0.005 | 4 | 8 |
| Per field potentials (group II) | 85 ± 10 |  |  | 97 ± 8 | 6 | -12% | 0.002 |  |  | 5 | 6 |

However, the quantitative comparison of the effects of peripheral conditioning has been hampered by the variability of these effects depending on experimental conditions. They depended in particular on the size of the test responses. This is illustrated in Figure 3e with a range of effects of the same intensity of the conditioning stimulation (Tib 4T) on field potentials of different sizes (evoked by graded epidural stimulation). For these reasons, the individual quantitative comparisons illustrated in Figures 3-6 were made on the effects of conditioning stimuli on test responses of matching sizes. The comparison summarized in Table 1 was also restricted to the effects of a standard intensity of conditioning stimuli (4-5T) on responses evoked by the optimal intensity test stimuli. Nonparametric Kruskal-Wallis test with Dunn’s multiple comparisons did not reveal any statistical differences (*p* > 0.05) between changes in field potentials and nerve volleys at any intensities of the conditioning stimuli (<2, 2, 3, 4, 5, 10T), although in individual experiments differences in both directions were observed (as indicated in Figure 3a,b).

**FIGURE 3.**
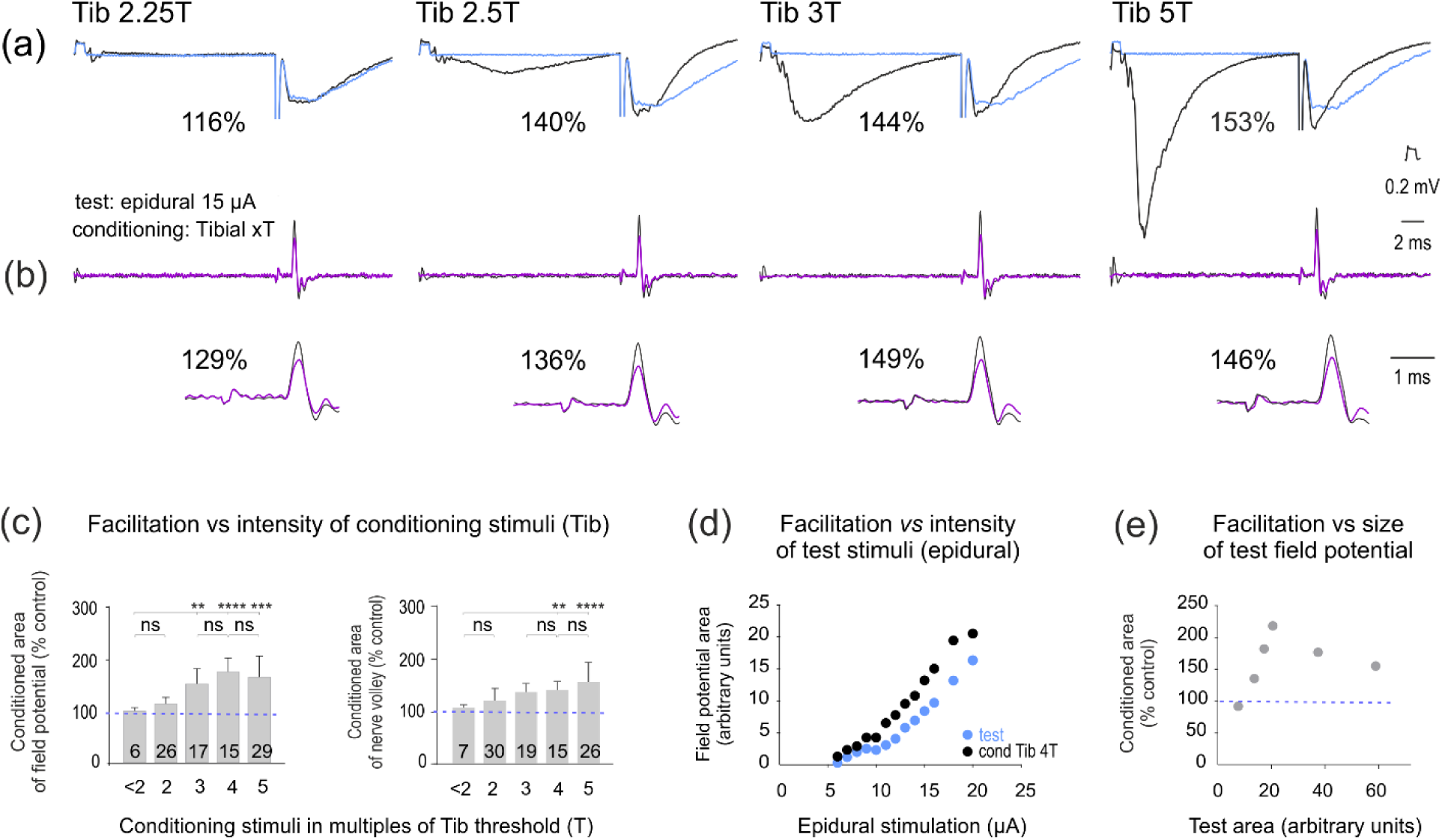
Facilitation of nerve volleys and the earliest components of field potentials evoked by epidural stimulation, reflecting the excitability of group I and group II muscle afferents, following the preceding conditioning stimulation of group II afferents in the tibial nerve (as outlined in Figure 1d). (a) Examples of dorsal horn field potentials evoked by 15 µA epidural stimulation preceded by conditioning stimulation of Tib at the indicated intensities. (b) Nerve volleys simultaneously recorded from Per, at the same time base as the field potentials and horizontally expanded. Superimposed records of control (blue and purple) and conditioned (black) responses with shock artefacts of epidural polarization truncated. Note the increase of the conditioned responses in % of control, indicated to the left, with the increase in the intensity of the conditioning stimuli. Note also that the increase in the early components of the field potentials was combined with the decrease in the later components attributable to presynaptic inhibition. For more details see Figure 4. (c) Mean increases in the areas of the early components of field potentials and nerve volleys evoked by a medium-intensity epidural stimulation preceded by conditioning stimulation of Tib at increasing intensities. Mean (± SD) increases in percentages of the control test responses. The number of trials in the samples is indicated on the bars. (d) An illustration of the relationship between the degree of increase in the area of the field potentials (in arbitrary units) and the intensity of epidural stimulation at a constant intensity of the conditioning stimuli (Tib 4T) in one of the experiments. ( e) An example of different degrees of facilitation of responses of different sizes, similarly evoked at a constant conditioning stimulus intensity (Tib 4T). The nonparametric Kruskal-Wallis test revealed a significant difference in medians among the 5 groups of data related to the intensity of conditioning stimuli (<2, 2, 3, 4, 5T) regarding the field potential as well as afferent volleys (H statistics significant at *p* < 0.0001). Further post-hoc analysis using Dunn’s test for multiple comparisons indicated statistically significant differences for conditioned responses as indicated by ** *p* < 0.01; *** *p* < 0. 001; **** *p* < 0.0001.

Taken together, these results indicate that group II but not group I muscle afferents, contribute to the modulation of input to the spinal cord from muscle afferents and that their effects resemble those of epidural polarization (Jankowska & Hammar, 2021), as well as previously reported effects of stimulation of unspecified peripheral afferents (Bączyk & Jankowska, 2018); for the differences, see Discussion. They also show that the preceding stimulation of group II afferents increases the excitability of both group I and group II afferents, thereby providing an important clue for the conclusions on the most likely relays of these facilitatory effects (see Discussion).

### Effects of the astrocyte toxin L-AAA on changes in the excitability of group I and II muscle afferent fibres at their entry to the spinal cord evoked by peripheral conditioning stimulation

Strong indications that the long-lasting effects of epidural polarization are mediated by astrocytes have been provided by showing that the astrocyte toxin L-AAA counteracts these effects (Hammar & Jankowska, 2024). We have now addressed the question as to whether astrocytes might likewise mediate the increase in the excitability of afferent fibres by conditioning stimulation of peripheral afferents. To this end, we compared the effects of conditioning stimulation of Tib as described above, with the effects after application of L-AAA.

In keeping with previous observations (Hammar & Jankowska, 2024), the excitability of epidurally stimulated fibres changed only marginally during the diffusion of L-AAA from a pipette in contact with the surface of the spinal cord. However, the effects of conditioning stimulation of Tib on the excitability of epidurally stimulated fibres were consistently weakened in the presence of L-AAA. As illustrated in Figure 4a,b,c and summarized in Table 1 (columns c,e,f), the effects of Tib during diffusion of L-AAA were weaker than under control conditions and the difference was highly statistically significant in all three experiments. The early components of epidurally evoked dorsal horn field potentials were then increased following Tib stimulation by 31% less than under control conditions (Table 1 column e). In one experiment in which no data prior to L-AAA diffusion were obtained but L-AAA was allowed to diffuse for 1 hour, the dorsal horn field potentials were increased by the conditioning Tib stimulation to only 104%. In addition, in one of the experiments, the latency of the dorsal horn field potentials was increased following conditioning Tib stimulation (from 0.7 to 1.09 ms from the stimulus, i.e. by 0.3-0.4 ms (Figure 4b), returning to the original latency about 2 hours after the diffusion of the toxin was terminated.

**FIGURE 4.**
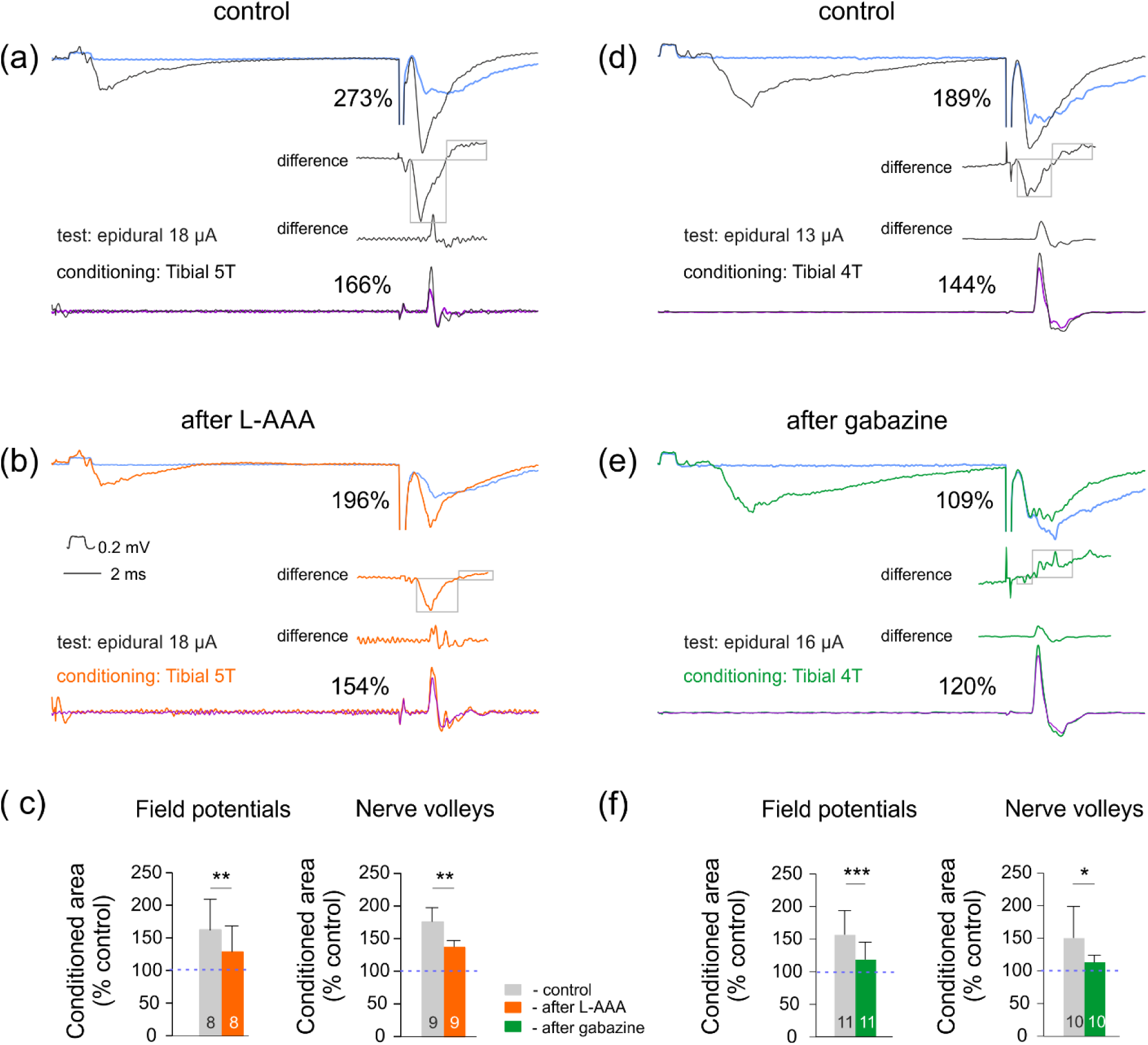
Effects of conditioning stimulation of a peripheral nerve (Tib) on the excitability of group I and II afferents excited by epidural stimulation under control conditions and under effects of the astrocyte toxin L-AAA the GABA channels blocker gabazine. (a) and (b) Upper traces, dorsal horn field potentials recorded at a depth of 0.85 mm. Superimposed records of effects of epidural stimulation (blue) and when it was preceded by conditioning stimulation of Tib under control conditions (black), and 42 min after the onset of administration of L- AAA (orange). The relative increases of the conditioned earliest components indicated to the left are in percentages of control values. The differences between the test and conditioned responses are below. Bottom traces, records from the peroneal nerve, the first components of the nerve volleys representing responses of group I afferents and later components responses of group II and/or undefined slower conducting afferents with the difference between the test and conditioned responses above them. (d) and ( e), as in (a) and (b) but for effects of gabazine tested 17 min after the onset of its administration in another experiment. Voltage calibration in b is for the intraspinal records. (c) and (f) Mean decrease in the facilitation of field potentials and nerve volleys under the effects of L-AAA and gabazine. * *p* <0.05,** *p* < 0.01 and *** *p* < 0.001 indicate statistically significant differences between the test and conditioned responses. See Methods and Figure 3.

### Effects of gabazine on changes in the excitability of group I and II muscle afferent fibres at their entry to the spinal cord evoked by peripheral conditioning stimulation

The increase in the excitability of epidurally stimulated fibres following stimulation of a peripheral nerve was considered to be due to field potentials evoked by such stimulation and to replicate effects of epidural polarization (Bączyk & Jankowska, 2018), with the silent assumption that the effects of stimulation of a peripheral nerve are likewise GABA-dependent. However, even if astrocytes rather than intraspinal field potentials contribute to the effects of a peripheral nerve the effects of peripheral and epidural stimulation are not necessarily mediated by the same categories of astrocytes. Recent studies revealed that astrocytes might increase the excitability of afferent fibres (as judged from their effects on afferent ganglion cells in the trigeminal nucleus by lowering [Ca2+] “at focal points along the axons” (Gaudel *et al*., 2022; Gaudel *et al*., 2024). Both in the trigeminal nucleus and in the visual cortex (Ryczko *et al*., 2021) the Ca 2+ related effects of glial cells modified the excitability of Nav 1.6 ion channels. Such astrocytes might thus also be involved in mediating effects of conditioning stimulation of Tib on afferents entering the spinal cord and act independently, or in parallel with GABAergic astrocytes activated by epidural polarization.

It thus became critical to test the effects of the GABA channel blocker gabazine on the facilitatory effects of Tib stimulation. Gabazine effectively counteracted increases in the excitability of epidurally stimulated afferent fibres during as well as following the period of their polarization (see Fig. 4e in Hammar & Jankowska, 2024). Provided that the effects of Tib stimulation are mediated by GABAergic astrocytes, they should likewise be reduced by gabazine.

The effects of gabazine were examined in the same way as the above-described effects of L-AAA and were indeed found to be similar (Table 1, columns b,d). Figure 4d,e,f illustrates the much weaker facilitatory effects of Tib stimulation during gabazine application compared to those evoked under control conditions. In the records illustrated in Figure 4, the early components of the dorsal horn field potentials evoked by epidural stimulation were increased to 109% during gabazine application rather than to 189 % in the control records. Similarly, the nerve volleys increased to 122% instead of 144 %. The mean differences amounted to 40% and 39% respectively (Table 1 column e).They were thus comparable to the mean differences in the presence of L-AAA and no statistically significant differences were found between the effects of gabazine and L-AAA on field potentials, although they were unexpectedly found with respect to nerve volleys.

### How long-lasting are the facilitatory effects of stimulation of group II afferent fibres?

Data in Figures 3-5 and in Table I show the effects of single conditioning stimuli applied to Tib about 20 ms before the test epidural stimuli when 10 test epidural stimuli (at 1 Hz) alternated with 10 such stimuli preceded by conditioning stimuli. Under such conditions, the mean increase of nerve volleys and field potentials fell within a range of 155-175% of the control (Table 1 column b) but was weaker than the increases following 1 min of epidural polarization (frequently exceeding 1000%). The duration of the effects was also shorter because the responses following the conditioning stimulus within a second range were usually unchanged. However, in some experiments, the successive responses were increased (up to 166%) when preceded by conditioning stimuli for 10-15 min, as illustrated in Figure 5a.

**FIGURE 5.**
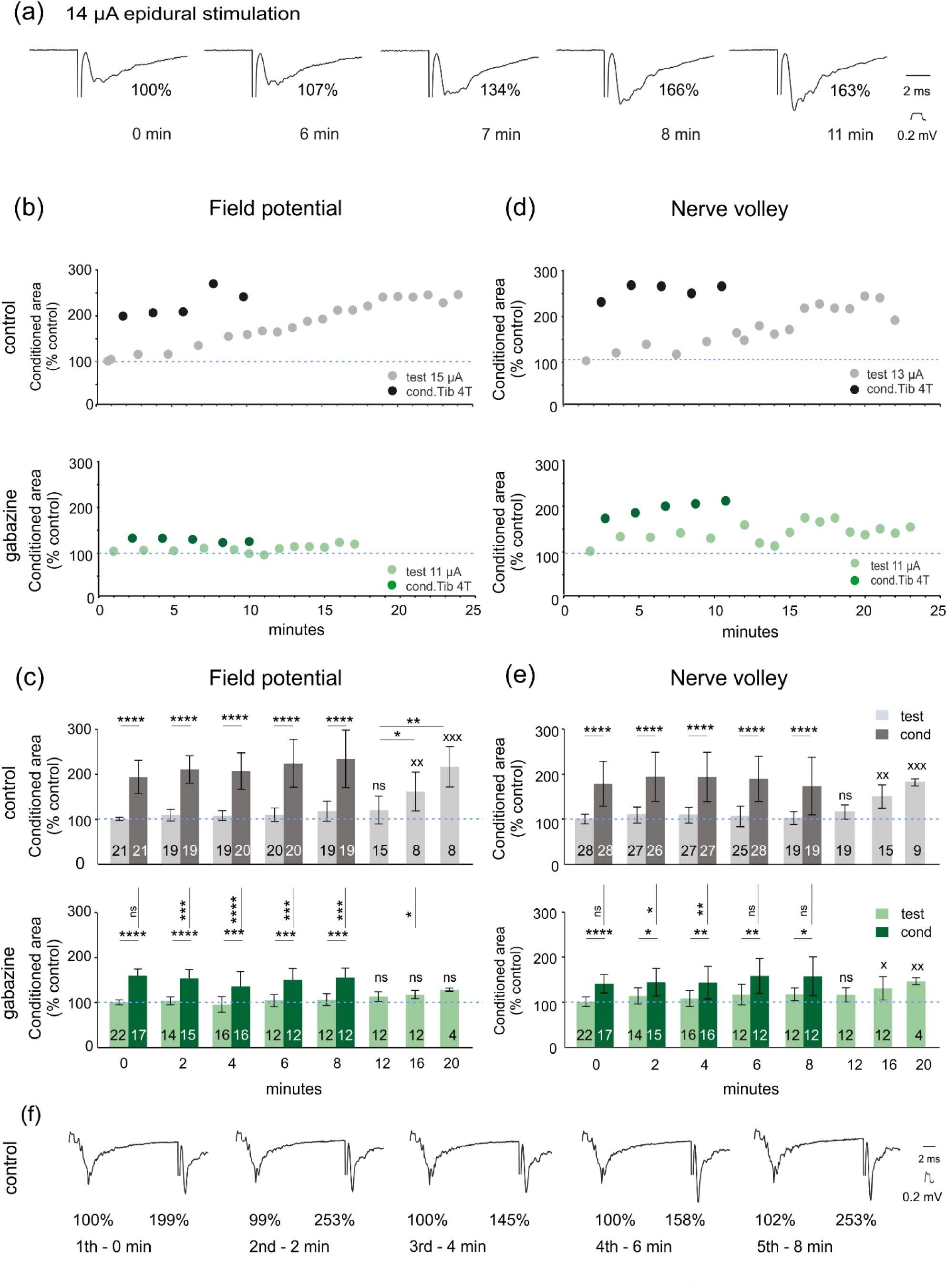
Prolonged facilitatory effects of peripheral afferent fibres. (a) Examples of field potentials evoked by epidural stimulation (14 µA, with shock artefacts truncated) in a 15-minute-long series of records when they were preceded by stimulation of Tib (3- 5T). Note an increase in the early components of these potentials. (b) and (d) Comparison of changes in 60 consecutive field potentials and nerve volleys evoked by epidural stimulation alone (grey) and when they were preceded by Tib stimulation (black) under control conditions and in the presence of gabazine (55 and 20 min from the beginning of its application, light and dark green). Data from two experiments with the largest effects on field potentials and nerve volleys respectively. Averages of 60 alternating test and conditioned responses during successive 1-minute periods and of 10 responses/min during the following 15 min. (c) and ( e) Mean decrease in the facilitation of field potentials and nerve volleys under effects of gabazine. (f) Comparison of field potentials evoked by Tib and epidural stimulation (left and right in each trace, with shock artefacts truncated, during successive 5 periods of conditioning stimulation in the series illustrated in (b) * *p* < 0.05, ** *p* < 0.01 and *** *p*<0.001, **** *p*<0.0001 indicate statistically significant differences between bars indicated by the horizontal lines (test *vs* conditioned responses) or the vertical lines (control conditioned responses vs conditioned responses under effects of gabazine). For the last three bars, ^x^ *p* < 0.05; ^xx^ *p*<0.01; ^xxx^ p < 0.001 and ns indicate the statistical significance of differences (or its absence) between responses evoked during the first control period and those following the 5^th^ conditioning sequence. See Methods and Figure 3. Note that the field potentials at 16 and 20 min (the last 2 bars) were significantly larger when compared to the first test ones but not after the gabazine application.

In order to maximize the effects of conditioning stimulation of peripheral afferents we modified the experimental conditions by using 5 trains of 60 test epidural stimuli at 1Hz alternating with trains of 60 epidural stimuli preceded by conditioning stimuli (Tib 4T). The increasingly stronger mean effects of conditioning stimuli applied in this way were found in 7/10 experiments. The nerve volleys increased within the range of 150% after the first series to 250% after the 3^rd^-4^th^ series and field potentials from 180% to 300% after the 4^th^-5^th^ series 10 min after the first control records (Figure 5b-e). In 3 experiments the nerve volleys and field potentials remained increased for 10-15 minutes following the fifth conditioning sequence, the two largest increases being illustrated in Fig. 5 b and d (top) but most often any post-conditioning increase did not exceed 120%.

In preparations in which L-AAA (1 experiment) or gabazine (4 experiments) were applied, the increases were much weaker. Plots illustrating the effects of gabazine on field potentials and nerve volleys in two of the experiments are shown in Figure 5b and d, with mean data being presented in Figure 5c and d. As indicated, all differences between responses evoked before and immediately following the trains of Tib stimulation (light and dark bars) were statistically significant as were also at least some changes between the responses evoked before the first sequence of conditioning stimuli and those evoked after the last of these sequences. The two plots also show statistically significant differences between the effects of conditioning stimulation under control conditions (dark grey bars) and those affected by gabazine (dark green bars).

Weaker effects of Tib conditioning stimuli in the presence of L-AAA and gabazine were associated with the previously reported shorter lasting effects of epidural polarization on the increase of both nerve volleys and dorsal horn field potentials (not illustrated) following the experimental design in Figure 1b.

The variability of effects of conditioning stimulation of a peripheral nerve evoked by either single stimuli or trains of stimuli at 1Hz lasting 1 minute appeared to be related to the changing background at which it was applied. Weaker effects were apparently evoked at the background of the excitability of the fibres modified by previously repeated stimuli and the largest effects of conditioning stimuli were seen at the beginning of the experiment. One of the factors to consider was the degree of depolarization of dorsal horn neurons by conditioning stimuli as this could affect the postsynaptic potentials reflected in extracellular field potentials induced in the dorsal horn neurons more than the number of the epidurally stimulated nerve fibres giving rise to them. It was therefore important to compare the effects of the conditioning Tib stimulation on dorsal horn field potentials evoked by epidural stimulation and by peripherally stimulated afferents. In the series of records illustrated in Figure 5f the field potentials following Tib stimulation remained practically unchanged (99-102%) while field potentials evoked by epidural stimulation increased (to 145-253%). Similar differences were seen in other experiments.

Taken together the reported effects of gabazine are in keeping with GABA-mediated effects of peripheral afferents both within seconds following their stimulation and for at least several minutes. Nevertheless, the effects of gabazine could only partly support the hypothesis of the involvement of the astrocytes in peripherally evoked modulatory effects because gabazine could counteract the effects of GABA released by GABAergic interneurons as well as by GABAergic astrocytes.

### Presynaptic inhibition of field potentials evoked by group II muscle afferents in preparations treated with astrocyte toxin L-AAA and gabazine

One of the factors interfering with the longer-lasting effects of peripheral afferents on the excitability of epidurally stimulated fibres might be presynaptic inhibition of transmission from the afferents (as outlined in Figure 1d). Under conditions of the present experiments, presynaptic inhibition might reduce synaptic actions of group II afferent fibres stimulated either peripherally or epidurally at least in the case of group II afferent fibres projecting to the dorsal horn. If the transmission between the epidurally stimulated fibres and their target neurons was weakened, the resulting field potentials would be reduced, and hence the numbers of these fibres, and estimates of their excitability deduced from the size of the field potentials might be underestimated. As shown in Figure 3e, both the smallest and near maximal dorsal horn field potentials evoked by epidural stimulation were less effectively increased than those within the middle range. The question therefore arose whether changes in presynaptic inhibition following L-AAA and gabazineadministration and the resulting decreases in the dorsal horn field potentials significantly interfered with the evaluation of the excitability of group II afferents.

To. address this question the origin of presynaptic inhibition within the explored region of the dorsal horn was first verified by comparing the degree of depression of field potentials evoked by Per group II afferents following conditioning stimulation of Tib. As illustrated in Figure 6a with records from one of the experiments, both early and later components of these potentials were decreased by Tib stimulation. The monosynaptic components of Per field potentials were on average decreased to 96-75% of control depending on the intensity of Tib stimulation (grey bars in Figure 6c). The degree of decrease of the later components could not be reliably quantified but appeared to be either similar or stronger. Both were decreased when Tib stimulation was within the range 2.5-5.0 times threshold while weaker conditioning stimuli were not effective and stronger stimuli failed to evoke increasingly stronger effects, in keeping with the origin of the presynaptic inhibition of transmission from group II muscle afferents (Rudomin & Schmidt, 1999).

**FIGURE 6.**
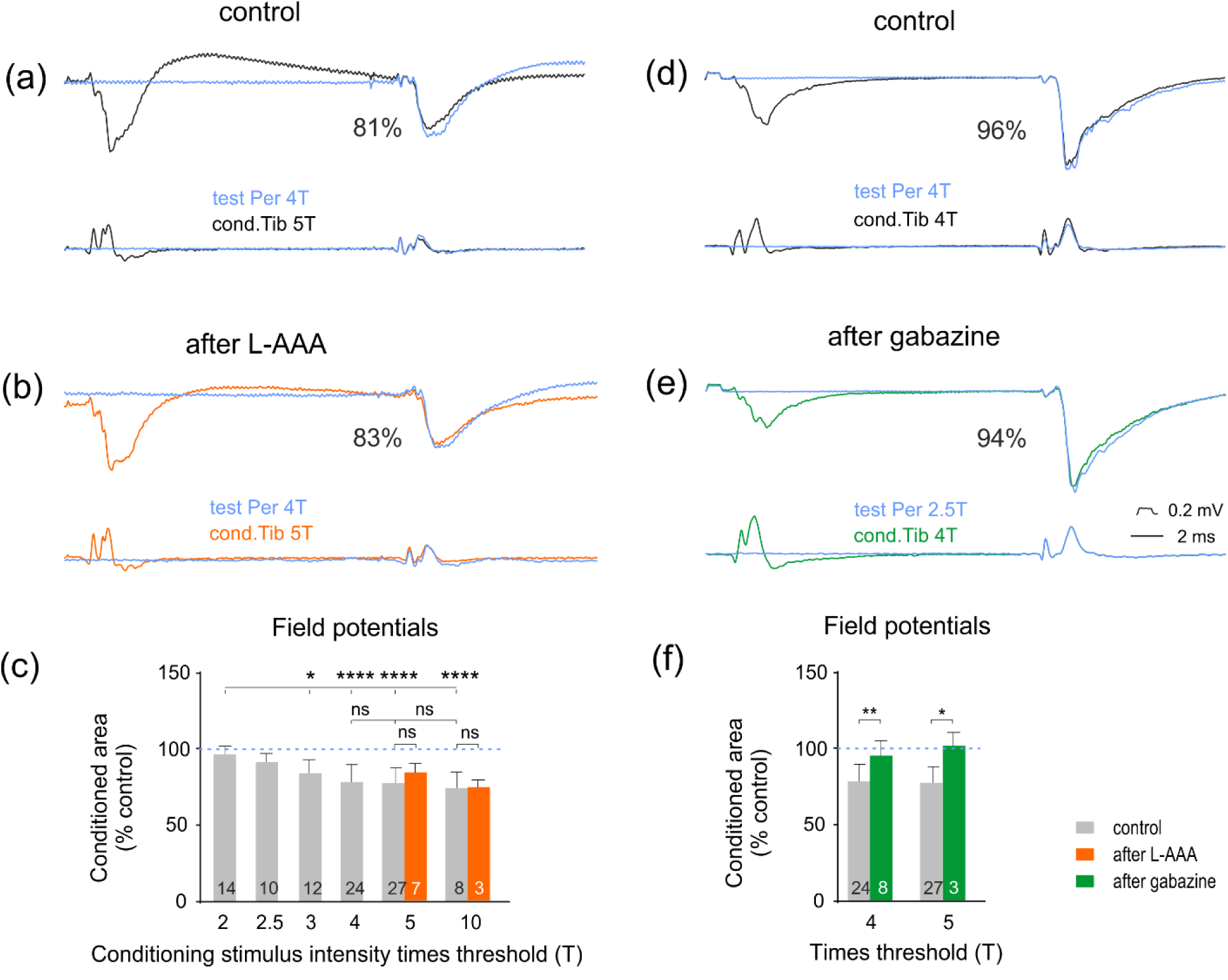
Comparison of effects of stimulation of a peripheral nerve on dorsal horn field potentials evoked by group II afferents in another nerve (presynaptic inhibition) under control conditions and in the presence of the astrocyte toxin L-AAA or the GABAA channel in the presence blocker gabazine. (a) and (b), Upper records, examples of field potentials evoked by stimulating Per alone (4T, blue traces) or preceded by stimulation of Tib (5T, under control conditions (black traces) and in the presence of the astrocyte toxin L-AAA (orange). Lower records, cord dorsum potentials. Changes in the areas of the earliest components of the conditioned field potentials are indicated to the left in percentages of control areas. (c) Mean areas of monosynaptic components of the test and conditioned field potentials (measured within a 0.9 ms time window) under control conditions and under effects of L-AAA. (d-f) as in (a-c) but for the effects of gabazine. * *p* < 0.05, ** *p* < 0.01 and **** *p*<0.0001 indicate statistically significant differences between the test and conditioned responses. See Methods and Figure 3. The nonparametric Kruskal-Wallis test revealed a lack of a significant difference between control and post-L-AAA conditions (H statistic *p* = 0.1173) while a statistically significant decrease in the presynaptic inhibition has been indicated by paired Student t-test in the presence of gabazine.

In L-AAA treated preparations the presynaptic inhibition of Per field potentials by Tib conditioning stimulation remained practically unchanged (Table 1 column c) or was only marginally decreased (Figure 6 a,b,c). Nevertheless, the paired t-test revealed that the statistical significance of differences between presynaptic inhibition before and during L-AAA treatment was close to its critical value (*p* = 0.54) not excluding the possibility that astrocytes might to some extent affect presynaptic inhibition and thus GABAergic interneurons that mediate it, albeit only marginally. In contrast, the decrease in the presynaptic inhibition by gabazine (from 85±10% to 97±8%, Table 1 column d) was statistically significant. To what extent the marginal decreases in presynaptic inhibition after L-AAA and the larger decrease by gabazine would affect estimates of the effects of Tib stimulation on the monosynaptic components of dorsal horn field potentials evoked by epidural stimulation would thus remain an open question. They might be in keeping with the slightly stronger relative increases of the facilitation of dorsal horn field potentials by Tib stimulation in the presence of gabazine than L-AAA (Table 1 column b).

However, they would not be in keeping with similar effects of Tib stimulation on dorsal horn field potentials and nerve volleys evoked by epidural stimulation which would not be expected to be affected by presynaptic inhibition. They should thus only have a minor impact on the estimates of the effects of the conditioning stimulation of peripheral afferents on the excitability of group II afferent fibres. Marginal effects of L-AAA on presynaptic inhibition might also indicate that astrocytes, in contrast to GABAergic interneurons may only have negligible effects on terminals of primary afferents.

## DISCUSSION

The results of this study are in support of the hypothesis that GABAergic astrocytes modulate spinal afferent input not only following epidural polarization but also when the modulation is evoked via spinal reflex pathways. The main argument in favour of this possibility is the demonstration that the astrocyte toxin (L-AAA) counteracts increases in the excitability of epidurally stimulated afferent fibres following conditioning stimulation of a peripheral nerve and that the GABAA channel blocker, gabazine, has a similar effect. An alternative possibility, that the increases in excitability are mediated by GABAergic interneurons has been found to be less likely; the arguments for and against the two options will be addressed in the second and third sections of the Discussion.

### Comparison of effects of conditioning stimulation of a peripheral nerve on the excitability of group I and II muscle afferents

Epidural stimulation may excite any nerve fibres within its effective radius. In the previous studies, we focused on changes in the excitability of group I muscle afferents, as judged from changes in the fastest conducting fraction of the fibres in which compound action potentials were evoked in a peripheral muscle nerve. Observations on other categories of epidurally stimulated afferent fibres were made only occasionally by noticing changes in later components of nerve volleys recorded in muscle nerves or skin nerves or in dorsal horn field potentials evoked by unspecified afferent fibres (Bączyk & Jankowska, 2018). However, at distinct locations within the dorsal horn monosynaptic field potentials following stimulation of a muscle or mixed peripheral nerves are evoked from specific afferents. In particular, in Rexed’s laminae III and IV such potentials are evoked by group II but not by group I afferents which have their target cells in more ventral laminae (Ishizuka *et al*., 1979; see their Fig. 9). Such field potentials may thus be attributed to either group II muscle afferents or the fastest conducting skin afferents, but to group II muscle afferents when evoked by stimuli suprathreshold for these afferents and at a latency required for group II mediated synaptic actions. In the present study, we evaluated, as previously (Jankowska & Hammar, 2021), the excitability of epidurally stimulated group I muscle afferents, from the earliest components of nerve volleys recorded in a peripheral nerve. The excitability of concurrently epidurally stimulated group II muscle afferents was estimated from changes in field potentials recorded at locations where they were only evoked at peripheral nerve stimulation intensities 2- 5 times threshold for group I afferents and at latencies 1.5-2.5 ms from group I afferent volleys but about 0.7 ms from group II afferent volleys. These latencies were in keeping with 0.6-0.8 ms longer conduction time along group II than group I muscle afferents (see Edgley & Jankowska, 1987b; in the cat, Riddell & Hadian, 1998; 2000 in the rat) and with one synaptic delay of 0.7-1.0 ms. We wish to stress that the classification of the explored dorsal horn field potentials as being of group II origin was based on the specificity of projections of afferent fibres to the selected area of the dorsal horn and not on the intensity of the epidural stimulation. As indicated in the methods section, the threshold epidural stimuli were the same for group I afferents giving rise to peripheral volleys and group II afferents evoking dorsal horn field potentials. Under our experimental conditions, the thresholds for activation of different fibre categories would thus depend on the distance from the tip of the stimulating electrode rather than the diameter of the fibres which is a decisive factor when they are stimulated within the peripheral nerves (size principle of Henneman, 1957). The epidural thresholds for both group I and II afferents were usually 10-12 µA and the stimuli used to evoke test responses were 16-18 µA.

The contribution of skin afferents to the explored field potentials could not be estimated. It was possible in view of the convergence of group II and skin afferents on neurons within the feline dorsal horn (Edgley & Jankowska, 1987b), although primarily within its medial part in the mouse, as judged from the projection area of skin and muscle afferents in this species (Gradwell *et al*., 2024) and the location of interneurons processing information from skin afferents (Hongo et al., 1989; Koch et al., 2018; Schouenborg 2002). Skin afferents did not nevertheless appear to contribute in a major way to the field potentials at the explored locations because these potentials were evoked at stimulus intensities exceeding 1.8 - 2T and such stimuli would excite a considerable proportion of skin afferents in a mixed nerve.

Based on these premises, the changes in peripherally recorded nerve volleys and in the first components of dorsal horn field potentials were used as a measure of changes in the excitability of epidurally stimulated group I and II afferent fibres respectively. The results revealed similar effects of the peripheral conditioning stimulation on group I and group II fibres (Figure 3, Table 1). The most effective conditioning stimulation of Tib was at intensities 2.5 to 4T. As no effects were evoked at conditioning stimulus intensities <2T, the contribution of group I muscle afferents and of low threshold skin afferents in Tib was unlikely and group II muscle spindle afferents appeared to play a major role. Furthermore, no major differences were found when conditioning stimulation of Tib was followed by small or large field potentials from this nerve within the explored region of the spinal cord, hence weakening the previously considered possibility of decisive effects of synaptically evoked electric fields.

Single 2-5T stimuli were followed by an increase in the excitability of epidurally stimulated afferent fibres up to 200-300%, i.e. weaker than that evoked by epidural polarization. However, the increase might have been underestimated if field potentials evoked in the dorsal horn were at the same time counteracted by presynaptic inhibition. As illustrated in Figures 3 and 6, an increase in the early components of these potentials by group II afferents in the tibial nerve was consistently combined with the decrease in the later components in parallel with presynaptic inhibition of peripherally evoked field potentials (Figures 5 and 6). The increase in the early components might thus have been evoked at the background of presynaptic inhibition. However, presynaptic inhibition or post-activation depression should not counteract the longer-term effects of stimulation of peripheral afferent fibres illustrated in Figure 5 and these were similar on group I and II afferents.

The presently reported effects of peripheral conditioning were found to be shorter lasting than the effects of epidural polarization but facilitatory effects of peripheral conditioning lasting at least one hour have been previously reported (Bączyk & Jankowska, 2018). The duration of effects of peripheral conditioning may however depend on experimental conditions, e.g. the distance between the sites of epidural stimulation with respect to the projection area of epidurally stimulated fibres. Considering the major role of intraspinal electric field potentials, Bączyk and Jankowska (2018) might have applied the epidural stimulation more caudally than in the present experiments to be closer to the region where Tib stimulation evoked larger dorsal horn field potentials, which were in addition evoked by weaker stimuli (2T). The effectiveness of 2T Tib stimuli might also indicate a stronger contribution of more caudally projecting skin afferents co-excited with group II afferents.

In summary, this comparison leads to the conclusion that the conditioning stimulation of group II afferents has a similar effect on the excitability of group I and II afferents and that the increase in their excitability by peripheral conditioning might be weaker, but not much weaker than by epidural polarization. These similarities might thus indicate that epidural polarization and stimulation of peripheral afferents modulate the excitability of group I and II afferents at the same site, likely at their branching region at the border between the dorsal column and the dorsal horn, and that these effects are mediated in a similar way.

### Contribution of astrocytes to the increase in the excitability of afferent fibres following stimulation of a peripheral nerve

The long-lasting increase in the excitability of afferent fibres at the border between the dorsal column and the dorsal horn following epidural polarization was originally attributed to direct effects of polarization of these fibres (Li *et al*., 2020). A similar increase in the excitability of dorsal column fibres following stimulation of a peripheral nerve (Bączyk & Jankowska, 2018) was therefore considered as due to extracellular field potentials evoked by peripheral afferent fibres projecting to the dorsal horn.

However, the sustained GABA-dependent increase in the excitability of afferent fibres by direct effects of epidural polarization has since been found to be negligible because it was prevented in the presence of the GABAA receptor blocker gabazine (see Fig. 4e in Hammar & Jankowska, 2024). The depolarization of GABAergic astrocytes by epidurally applied DC and the resulting increase in the concentration of GABA released by them appeared to be critical because the long-lasting effects of epidural polarization were prevented by the astrocyte toxin L-AAA (Hammar & Jankowska, 2024) as effectively as by gabazine. It was therefore considered that extracellular dorsal horn field potentials evoked by primary afferent fibres might likewise have a negligible direct effect on branching regions of afferent fibres but contribute to the depolarization of the astrocytes and their GABA-mediated effects. Astrocytes might also be excited by glutamate spillover from terminals of peripheral afferents terminating on dorsal horn neurons or of terminals of these neurons targetting other dorsal horn neurons. In both cases, the GABA channel blocker acting within the postulated region of actions of astrocytes should counteract the effects of the conditioning stimulation of peripheral afferents, as indeed found. The results illustrated in Figures 4-6 show that gabazine effectively counteracted the Tib-evoked increases in the excitability of afferent fibres. The hypothesis that these GABA-mediated effects are mediated by astrocytes is further supported by the finding that L-AAA weakened these modulatory effects. However, as summarized in Table 1, L- AAA counteracted peripherally evoked modulation in the excitability only moderately. This might indicate a less critical contribution of astrocytes to the peripherally than to epidurally evoked modulation. However, the excitability of epidurally stimulated fibres during epidural polarization (i.e. the phasic effect of epidural polarization) was likewise only moderately decreased in the presence of L-AAA, (cf Fig. 4 in Hammar & Jankowska, 2024), Hence, relatively weak interaction with the long-lasting increase in the excitability of nerve fibres by peripheral afferents does not contradict the postulated mediation of modulatory effects of peripheral afferents by astrocytes.

The relatively weak effects of L-AAA might be related to several factors. For instance, only a relatively small proportion of the dorsal horn astrocytes was found to be GABAergic (Christensen *et al*., 2018), their distribution might not be uniform and different subpopulations of GABAergic astrocytes might be affected by epidural polarization and peripheral conditioning stimulation. Those excited by low threshold skin and group II muscle afferents, constituting 11.29% of the total astrocyte population in laminae III-V according to Xu *et al*. (2021), might in addition be located more ventrally. Different subpopulations of astrocytes might also be affected by glutamate and GABA. In addition, the variability of the effects of L-AAA may depend not only on space but also on the timing of our tests with respect to the beginning of the administration of L-AAA. According to Khurgel *et al*. (1996), L-AAA causes ablation of astrocytes within the region of its injection within 48 hours, although immunohistochemical changes occur within about an hour and functional effects of L-AAA in vivo were noted within 0.5-1 hour (Chang *et al*., 1997; Xu *et al*., 2021).

Some of the requirements for GABA-mediated effects of peripherally stimulated afferents have already been satisfied. In particular, the study by Christensen *et al*. 2018) demonstrated that astrocytes may release GABA upon an increase in the concentration of glutamate released by afferent fibres, or by neurons excited by them. Astrocytes might have also contributed to the reported increase in fibre excitability in other ways, e.g. by modulating the concentration of calcium in the extracellular space within the branching region of these fibres at their entry to the spinal grey matter, as in the trigeminal nucleus (Gaudel *et al*, 2024) or in the visual cortex (Ryczko *et al*., 2021). Astrocytes constitute a highly non-homogeneous population (see e.g. Khakh & Sofroniew, 2015; Hanani & Verkhratsky, 2021) so that their subpopulations in different regions of the brain and the spinal cord could be involved in either different or similar ways. However, astrocytes could be affected by glutamate spilt at the site of their location, whether by group II and/or skin afferents in the dorsal horn of the spinal cord or its other sources in the trigeminal nucleus or the visual cortex.

### An alternative to the major contribution of astrocytes to the increase in the excitability of afferent fibres following stimulation of a peripheral nerve

Stimulation of peripheral afferents might evoke an increase in the excitability of epidurally stimulated afferent fibres via GABAergic interneurons as well as astrocytes. However, as the effects of PAD are short-lasting and do not exceed a fraction of a second, any longer-lasting effects of conditioning stimulation of peripheral afferents on either group I or II muscle afferents would be difficult to attribute to GABAergic interneurons by themselves. Furthermore, the contribution of GABAergic interneurons with input from group II and skin afferent fibres would be in keeping with the origin of PAD of group II muscle afferents but not of group I afferents (for references see Rudomin & Schmidt, 1999). GABAergic interneurons would thus not be likely to account for changes in the excitability of group I afferents by conditioning stimulation of group II afferents.

The major contribution of GABAergic interneurons to the increase in the excitability of group II afferents might also be questionable in view of the opposite effects of Tib conditioning stimulation on the early (monosynaptic) and later (disynaptic) components of the dorsal horn field potentials evoked by epidural stimulation (Figures 4 and 6). Both effects are evoked by group II afferents but while the early components are increased the later components are decreased. If the increase were mediated by astrocytes and the decrease by GABAergic interneurons reflecting presynaptic inhibition and PAD evoked via axo-axonic contacts with group II and skin afferents terminating within the dorsal horn, the explanation would be fairly simple. It would also be in keeping with the differences in the effects of spinal injuries on GABA receptors on afferent fibres, which increase in parallel with the increased spinal excitability, while the number of terminals of GAD2 interneurons remains unchanged (Hari *et al*. 2022, 2024). If the source of GABA affecting afferent fibres were restricted to GABAergic interneurons, distinct subpopulations of such interneurons would have to be postulated and the hypothetical interneurons affecting the excitability of the epidurally stimulated group I but not presynaptic inhibition of transmission from these afferents nor their PAD would have to be specified.

Using molecular features of neurons expressing *Slc32a1* (vesicular GABA transporter, Vgat), Haring Haring *et al*. (2018) identified 15 distinct clusters of GABAergic neurons within the dorsal horn, postulating at least 15 GABAergic neuronal types in addition to any other populations located more ventrally. Several of these might contribute to the GABA-mediated modulation of fibre excitability. The distribution of some of these 15 clusters corresponded to the distribution of GABAergic neurons in genetically modified mice in which PAD was evoked optogenetically, both in vivo and under conditions of acute experiments in vitro and were reported to be activated via V3 neurons (Lin *et al*., 2023). At an apparently similar location in the medial part of laminae V and VI of Rexed were also found inhibitory interneurons distinguished as RORbeta interneurons (Koch et al., 2017) and concluded to contribute to presynaptic inhibition via GAD2-expressing terminals in contact with primary afferents. Which of PAD- mediating GABAergic interneurons might be involved in addition to, or instead of the astrocytes in the reported modulation of fibre excitability is an open question. As reviewed by Rudomin & Schmidt (1999) different subpopulations of GABAergic interneurons are likely to contribute to presynaptic inhibition and/or primary afferent depolarization (PAD) of different categories of afferent fibres, in particular group I on the one hand and group II and cutaneous afferents on the other. GABAergic interneurons targeting group I and group II afferents should thus be excited by different subpopulations of V3 interneurons and/or RORbeta interneurons, most likely located in the intermediate zone (with group Ia and Ib input) or laminae III and IV interneurons (with group II and cutaneous monosynaptic input) respectively (see Riddell *et al*., 1995). However, modulation of fibre excitability might also be mediated by GABAergic interneuronsnot involved in presynaptic inhibition or even by glutamatergic neurons with undefined patterns of input (Rudomin, 2000; Russo *et al*., 2000; Lin *et al*., 2023). If some glutamatergic interneurons with group II input evoke previously undisclosed depolarization of group I afferent fibres (Lin *et al*., 2023), such hypothetical interneurons might also contribute to facilitatory effects of peripheral afferents on group I afferents together with those evoking PAD of group II afferents. The reported modulation of the excitability of afferent fibres at the level of their entry to the spinal cord might thus be explained, by assuming joint actions of astrocytes and various combinations of spinal interneurons.

### Some functional consequences

By modulating input from primary afferents, astrocytes may set up the level of excitability of spinal neuronal networks by natural stimuli as a part of feedback or feed-forward adjustments under different behavioural conditions. To comply with this, the population of astrocytes would need to be differentiated, and it has been repeatedly stressed that it is a highly heterogeneous population. In the context of the present study, it is of interest that only a subpopulation of astrocytes in the dorsal horn has been found to be GABAergic (Christensen *et al*., 2018), i.e. could directly modulate the excitability of afferent fibres entering the spinal grey matter. It is also relevant that astrocytes at different locations may modulate neuronal excitability and synaptic transmission in different neuronal circuits (for references see Xu *et al*., 2021). Astrocytes activated by group II muscle afferents should thus be among the laminae III-IV subpopulation. Whether they belong to the GABAergic astrocytes and what are their spinal targets has not been specified.

It is also of importance that the postulated facilitatory actions of astrocytes activated by muscle stretches or by a sufficiently weak spinal stimulation would not be at variance with inhibitory actions of more dorsally located astrocytes depressing synaptic transmission in pain pathways from nociceptors. By activating different sub-populations of astrocytes, it should thus be possible to take advantage of their different modulatory actions to the benefit of the patients.

How the modulatory actions of astrocytes are combined with unspecific excitatory effects evoked via GABAergic interneurons and more specific effects of PAD remains an open question.

## Abbreviations and terminology

ECG,: electrocardiogram
GABA: gamma-aminobutyric acid
GAD2,: glutamate decarboxylase 2 enzyme for GABA biosynthesis
L-AAA: astrocyte toxin L-alpha-aminoadipic acid
L3-L4: 3^rd^ and 4^th^ lumbar spinal cord segments
PAD: primary afferent depolarization
Per: common peroneal nerve
T,: the threshold for group I afferent stimulation
Tib: tibial nerve
Epidural polarization: polarization by constant direct current
Epidural stimulation: stimulation by constant intensity current pulses

## Acknowledgements

We wish to thank Drs D. Bennett and M. Bączyk for their helpful comments on the earlier versions of the manuscript. The study was supported by the Institute of Neuroscience and Physiology, Göteborg University (for IH and EJ), The Royal Society of Arts and Sciences (Kungl. Vetenskaps och Vitterhets-Samhället KVVS) in Göteborg (for EJ) and Nencki Institute of Experimental Biology (for US).

## Authors contribution

All authors contributed to the study design, data collection, interpretation and drafting of the manuscript. Data analysis and preparation of the figures by US and EJ.

